# Casein kinase II promotes piRNA production through direct phosphorylation of USTC component TOFU-4

**DOI:** 10.1101/2023.08.09.552615

**Authors:** Gangming Zhang, Chunwei Zheng, Yue-he Ding, Craig Mello

## Abstract

Piwi-interacting RNAs (piRNAs) are genomically encoded small RNAs that engage Piwi Argonaute proteins to direct mRNA surveillance and transposon silencing. Despite advances in understanding piRNA pathways and functions, how the production of piRNA is regulated remains elusive. Here, using a genetic screen, we identify casein kinase II (CK2) as a factor required for piRNA pathway function. We show that CK2 is required for the localization of PRG-1 and for the proper localization of several factors that comprise the ‘upstream sequence transcription complex’ (USTC), which is required for piRNA transcription. Loss of CK2 impairs piRNA levels suggesting that CK2 promotes USTC function. We identify the USTC component twenty-one-U fouled-up 4 (TOFU-4) as a direct substrate for CK2. Our findings suggest that phosphorylation of TOFU-4 by CK2 promotes the assembly of USTC and piRNA transcription. Notably, during the aging process, CK2 activity declines, resulting in the disassembly of USTC, decreased piRNA production, and defects in piRNA-mediated gene silencing, including transposons silencing. These findings highlight the significance of posttranslational modification in regulating piRNA biogenesis and its implications for the aging process. Overall, our study provides compelling evidence for the involvement of a posttranslational modification mechanism in the regulation of piRNA biogenesis.

## Introduction

Piwi-interacting RNAs (piRNAs) associate with Piwi-clade Argonautes in metazoans ^1, 2^. piRNAs regulate diverse biological processes including virus defense ^3^, sex determination ^4^, male fertility ^5^ and genome stability ^2^. In *C. elegans*, piRNAs engage thousands of germline-expressed mRNAs ^6^ and are required for recognizing and silencing newly-introduced foreign transgenes ^7^.

In *C. elegans*, piRNAs are also known as 21 U-RNAs. 21 U-RNAs are transcribed by RNA polymerase II (Pol II) from thousands of genomic locations ^8^. There are generally two types of 21 U-RNA loci, type 1 loci appear dedicated to the production of piRNAs and are located in huge clusters on chromosome IV (LGIV), while type 2 piRNAs are produced often bidirectionally at most pol II transcription start sites, including those producing longer transcripts such as mRNAs or ncRNAs. Both types of piRNAs are transcribed as capped short RNAs (piRNA precursors) of approximately 27 nts in length. Type 1 piRNA genes contain an 8 nt sequence element called the Ruby motif ^9^ located approximately 40 nts upstream of their transcription start sites. Expression of type 1 (but not type 2) piRNA precursors depends on components of the ‘upstream sequence transcription complex (USTC)’—including PRDE-1, SNPC-4, TOFU-4 and TOFU-5 ^10^. After transcription piRNA precursors are trimmed by the PETISCO/PICS complex which is thought to remove the cap and first two nucleotides to expose an internal 5’ uridine residue necessary for stable association with Piwi Argonaute ^11, 12^. The conserved exonuclease PARN-1 trims piRNA 3’ ends from the piRNA precursors to generate the mature 21 nt piRNAs ^13^.

Casein kinase II (CK2) is a conserved serine/threonine kinase involved in many cellular processes including transcriptional regulation, DNA repair and cell survival ^14, 15^. Phosphorylation mediated by CK2 affects protein-protein interaction, for example, the phosphorylation of Rap1 by CK2 promotes Rap1/Bqt4 interactions to facilitate telomere protein complex formation ^16^.

Aging in animals is associated with extensive functional decline, ranging from a decline in tissue integrity, motility, learning and memory, and immunity ^17^. These changes are correlated with and likely caused at least partially by remodeling of the epigenome. However, the mechanisms underlying the loss of heterochromatin during the aging process remain largely unknown. Investigating the factors and mechanisms that contribute to the age-related loss of heterochromatin will provide valuable insights into the molecular basis of aging and its associated functional decline.

Here we show that CK2 is required for piRNA mediated silencing in *C. elegans*. Depletion of CK2 affects piRNA biogenesis. Furthermore, we demonstrate that CK2 is required for phosphorylation of the USTC component TOFU-4, which was previously reported to promote piRNA transcription ^10^. We observed that this phosphorylation process is impaired during the aging process, resulting in disruptions to piRNA-mediated gene silencing. Collectively, our findings strongly indicate that CK2-mediated phosphorylation of TOFU-4 plays a critical role in regulating USTC assembly, promoting piRNA transcription, and is required for effective piRNA-mediated gene silencing during the aging process.

## Results

### CK2 promotes piRNA mediated silencing

In *C. elegans* extensive forward genetic screens have identified genes required for piRNA silencing. However, a limitation of these previous studies is that they require the resulting strains to be both piRNA silencing defective and viable. We reasoned that many important regulators of piRNA biogenesis might also function in pathways essential to development and viability. To identify such factors, we performed an RNAi-based screen of genes previously determined to be essential for viability. To do this we used a sensor system expressing a bright easily scored GFP transgene whose silencing requires an active piRNA pathway (Fig. 1A) ^18^. In wild type worms, the sensor GFP signal is silenced within the pachytene zone of the ovary. However, in *Piwi* pathway mutants, bright GFP signal expands throughout the pachytene zone in the sensor animals (Fig. 1A, B).

**Figure 1.**
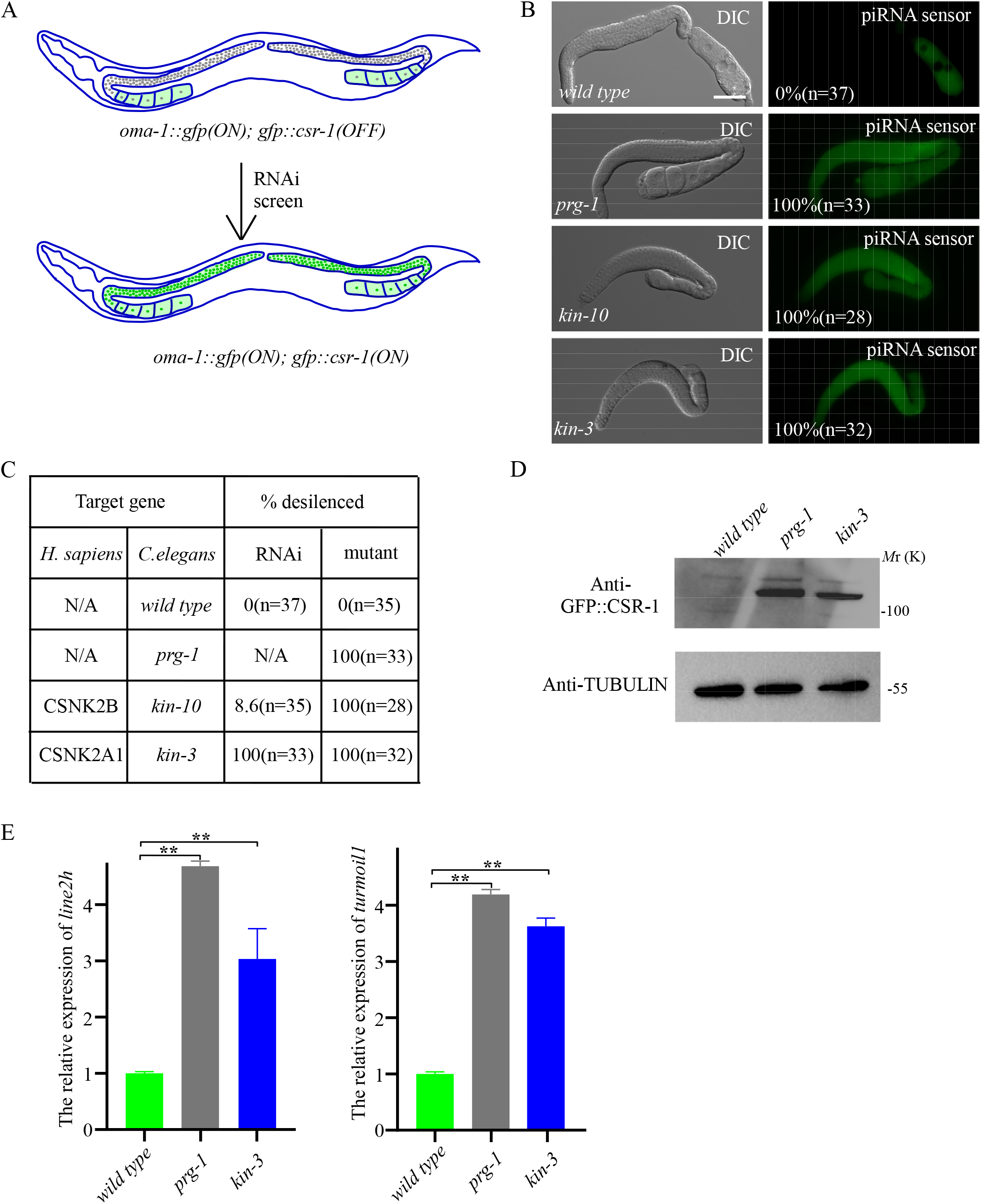
CK2 complex promotes piRNA mediated silencing. **(A)** Schematic overview of the piRNA sensor screen, the piRNA sensor contains an actively expressed OMA-1::GFP transgene and a GFP::CSR-1 transgene silenced by the piRNA pathway ^34^. Inactivation of piRNA pathway will result in the GFP::CSR-1 expression in the perinuclear region. **(B)** The GFP::CSR-1 transgene is silenced in wild type worms, while in *prg-1*, *kin-10* and *kin-3* mutants, GFP::CSR-1 is expressed. **(C)** Percentage of worms with expressed piRNA sensors in wild type, *prg-1*, *kin-10* and *kin-3* mutants. n. Total number of animals scored. **(D)** Western results of GFP::CSR-1 protein levels in wild type, *prg-1* and *kin-3* mutants. **(E)** qRT-PCR results of *line2h* and *turmoil* RNA levels in wild type, *prg-1* and *kin-3* mutants. mRNA level in wild type worms is set to 1.0. Data are shown as mean ± SD. **p < 0.01. Scale bars: 50 μm for B.

Sensor animals were exposed to RNAi by feeding groups of about 100 worms on petri plates seeded with individual bacterial strains each expressing a dsRNA targeting one of 945 essential genes. This screen identified 39 positives, including several pathways that were previously described as essential for both viability and piRNA silencing (Table 1). For example, the piRNA reporter was desilenced when *hda-1* and *smo-1* were depleted ^18^, as well as depletion of the integrator complex, which has been shown to terminate *C. elegans* piRNA transcription ^19^. Furthermore, we found that other genes, such as splicing factors and nuclear pore complex components, were also capable of desilencing the piRNA reporter. Of these genes, we were particularly intrigued to discover that knockdown of components of the Cassien Kinase 2 (CK2) complex led to desilencing of the piRNA sensor. For example, we found that RNAi of *kin-10* (a regulatory subunit) and of *kin-3* (a catalytic subunit) desilenced the piRNA sensor (Fig. 1C). To confirm the RNAi findings, we used an auxin-inducible degron system to conditionally deplete KIN-3 and KIN-10. Using CRISPR we inserted sequences encoding 46 amino acids corresponding to the plant degron into the endogenous *kin-3* and *kin-10* genes in a strain expressing the plant F-box protein TIR1 which upon auxin binding associates with the DEGRON domain to recruit the proteosome machinery ^20^. Exposure of this strain to auxin at 1mM completely desilenced the piRNA sensor (Fig. 1B-D), and upon prolonged exposure caused lethality as expected.

**Table 1.** Summary of RNAi-based genetic screen of lethal genes using a silenced piRNA sensor.

| RNAi | Score | Number | Description |
| --- | --- | --- | --- |
| L4440 | 0 | N=40 | Empty vector |
| *prp-19* | 43.75% | N=16 | splicing factor |
| *prp-17* | 100% | N=32 | splicing factor |
| *npp-1* | 29.41% | N=17 | Nuclear pore complex |
| *npp-8* | 25% | N=12 | Nuclear pore complex |
| *npp-6* | 44% | N=25 | Nuclear pore complex |
| *npp-7* | 45.45% | N=22 | Nuclear pore complex |
| *npp-11* | 14.29% | N=14 | Nuclear pore complex |
| *rpb-2* | 23.08% | N=13 | RNA Polymerase |
| *rpc-1* | 35.71% | N=14 | RNA Polymerase |
| *aco-2* | 22.22% | N=18 | mitochondrion protein |
| *cox-5A* | 15.38% | N=13 | mitochondrion protein |
| *dnj-10* | 26.32% | N=19 | mitochondrion protein |
| *mrps-22* | 11.76% | N=34 | mitochondrion protein |
| *ints-6* | 21.43% | N=14 | snRNA 3'-end processing |
| *spt-5* | 42.86% | N=14 | regulation of transcription |
| *spt-16* | 100% | N=20 | FACT complex |
| *xpo-1* | 16.67% | N=12 | nuclear export receptor |
| *xpo-2* | 90.36% | N=83 | nuclear export receptor |
| *smo-1* | 27.78% | N=54 | SUMO |
| *hda-1* | 25% | N=12 | NuRD complex |
| *let-92* | 21.05% | N=19 | Part of protein phosphatase type 2A complex |
| *gsp-1* | 41.18% | N=17 | protein serine/threonine phosphatase |
| *mcm-7* | 21.43% | N=14 | minichromosome maintenance complex |
| *mrg-1* | 83.64% | N=55 | chromodomain-containing protein |
| *trd-1* | 65% | N=20 | Tetratricopeptide repeat Regulator of Differentiation |
| *cpsf-2* | 7.69% | N=13 | mRNA 3'-end processing |
| *his-7* | 16.67% | N=12 | part of nucleosome. |
| *isy-1* | 23.08% | N=13 | ISY splicing factor homolog |
| *erfa-3* | 23.08% | N=13 | translation factor |
| *emb-4* | 29.41% | N=17 | mRNA splicing |
| *snpc-4* | 37.78% | N=45 | piRNA transcription |
| *use-1* | 78.95% | N=19 | vesicle transport |
| *C09H10.10* | 44.12% | N=34 | containing Forkhead-associate domain |
| *mog-5* | 40.26% | N=77 | DEAH helicase |
| *tgs-1* | 76.39% | N=72 | Trimethylguanosine synthase 1 |
| *C53H9.2* | 37.84% | N=45 | GTP binding protein |
| *ccm-3* | 26.09% | N=45 | programmed cell death protein 10 |
| *ncbp-1* | 20% | N=20 | nuclear cap-binding complex |
| *kin-10* | 8.57% | N=35 | protein kinase |

We next examined whether piRNA mediated silencing of transposable elements is disrupted in CK2 mutants ^21^. We examined the *line2h* and *turmoil1* transposons, which are transposons silenced by piRNA pathway. As expected, expression of the *line2h* and *turmoil1* transposon mRNAs were increased in *prg-1* and *kin-3* mutants (Fig. 1E). Together, these findings suggest that the CK2 complex promotes piRNA mediated silencing.

### CK2 promotes the levels of both mature and precursor piRNAs

We next wished to explore the effect of CK2 on piRNA levels. The levels of both mature and precursor piRNAs were determined as previously described ^22^ by deep sequencing small RNA populations from 2 biological replicates of *kin-3* degron animals after auxin exposure. For comparison we also analyzed small RNAs sequenced from wild type animals and from *prg-1* and *prde-1* mutants which are required respectively for stabilizing mature piRNAs and for promoting transcription of piRNA precursors. As previously shown ^22^, mature piRNAs were dramatically reduced compared to wild type levels in both *prg-1* and *prde-1* mutants (Fig. 2A, B) and were also dramatically reduced in *kin-3* depleted animals (Fig. 2A, B). Moreover, comparison of type 1 piRNAs in *kin-3* and *prg-1* mutants revealed similar levels of depletion across all ∼15,000 species (Fig. S1). Examination of type 2 piRNA levels revealed that *kin-3* caused at most a partial reduction in levels, an effect similar to that of *prde-1,* which is required for transcription of type 1 but not type 2 piRNAs (Fig. 2C)^22^.

**Figure 2.**
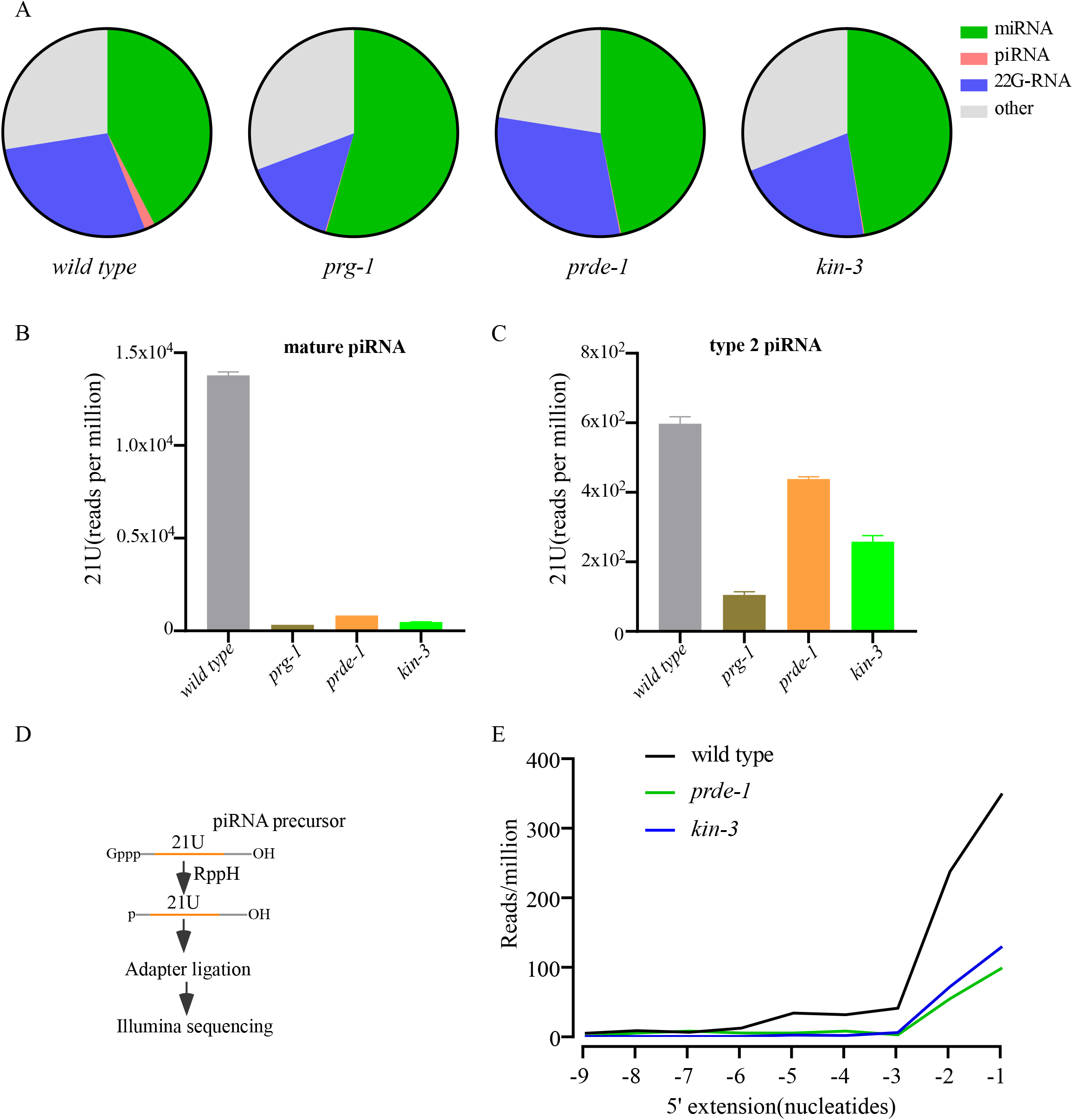
CK2 promotes the levels of both mature and precursor piRNAs. **(A)** The distribution of reads that correspond to genome annotation in small RNA libraries from wild type, *prg-1*, *prde-1*, and *kin-3* mutants was depicted using pie charts. **(B)** Bar diagram displaying piRNAs mapping perfectly to known *C. elegans* piRNA loci in wild type, *prg-1*, *prde-1* and *kin-3* mutants. **(C)** Bar diagram displaying type 2 21U-RNA abundance in wild type, *prg-1*, *prde-1*, and *kin-3* mutants. **(D)** Outline of strategy to detect the piRNA precursors by deep sequencing. **(E)** Distribution of 5’ extending nucleotides for the piRNA precursor sequences mapping to piRNA sequences. piRNA precursors from loci as above were examined.

Libraries prepared to enrich for the small capped RNAs that represent piRNA precursors revealed that piRNA precursor levels were reduced in the *kin-3* depleted libraries to levels similar to those found in *prde-1* mutants (Fig. 2D, E). Taken together, these findings suggest that CK2 is required for the production or stability of piRNA precursors.

### CK2 promotes the localization of USTC factors

Previous studies have shown that PRG-1 protein levels and its localization to peri-nuclear nuage, P granules, are both reduced in mutants that disrupt expression of piRNA precursors, while PRG-1 mRNA levels remain unaffected. These findings suggest that PRG-1 like some other Argonaute proteins such as Ago2 becomes unstable when unloaded ^23–25^. Consistent with these previous findings we found that in *kin-3* depleted animals, PRG-1 mRNA levels were not reduced compared to wild type (Fig. S2A), while in contrast, GFP::PRG-1 failed to localize in P-granules and instead localized diffusely in the cytosol of each mutant (Fig. 3A). We observed an identical change in GFP::PRG-1 localization in *tofu-4* (Fig. 3A), a factor previously shown to be required for piRNA transcription and for PRG-1 protein stability ^10^. When assayed by western blotting with GFP-specific antibodies in *kin-3* mutants and in *tofu-4* mutants, the band corresponding to full-length GFP::PRG-1 protein appeared strongly reduced and a 30KD band, likely representing free GFP, became prominent (Fig. 3B). As a control we monitored GFP::PGL-3, a P-granule marker, whose localization in nuage does not depend on piRNA biogensis (Fig. S2B, C), and found that GFP::PGL-3 is localized properly in the nuage of *kin-3* depleted animals.

**Figure 3.**
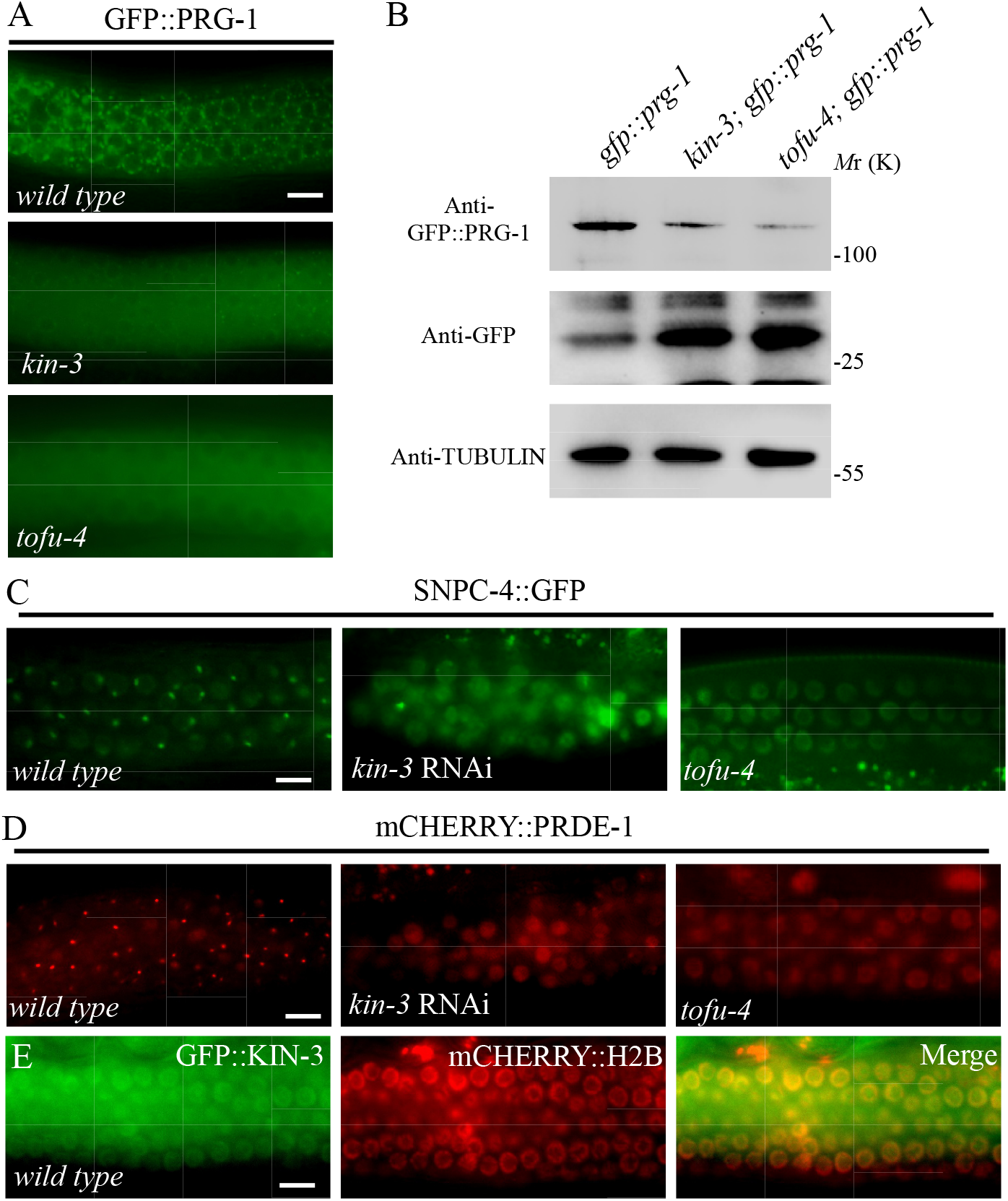
CK2 promotes the localization of USTC factors. **(A)** In wild type worms, GFP::PRG-1 is expressed in the perinuclear region, while in *kin-3* and *tofu-4* mutants, there is some diffused GFP signal. **(B)** Immunoblotting assays revealed that protein levels of GFP::PRG-1 are decreased while protein levels of free GFP is increased in extracts of *kin-3* and *tofu-4* mutants compared to those of wild type worms. **(C, D)** Compared to wild type worms, the SNPC-4::GFP (C) and mCHERRY::PRDE-1 (D) foci in *kin-3* and *tofu-4* mutants disappear. **(E)** GFP::KIN-3 in the nucleus is colocalized with the chromatin marker mCherry:H2B in young adult germ nuclei. Scale bars: 10 μm in A, C, D and E

SNPC-4 and PRDE-1 are two USTC factors required for piRNA biogenesis in *C. elegans* ^10^. In germline nuclei, they form foci that co-localize with the piRNA gene clusters on LGIV. We found that *kin-3*-depleted animals and as previously reported *tofu-4* mutants ^10^ failed to form subnuclear foci of SNPC-4::GFP and mCHERRY::PRDE-1 (Fig. 3C, D). These findings suggest that CK2 acts along with TOFU-4 to promote SNPC-4 and PRDE-1 localization to piRNA clusters where transcription occurs. GFP::KIN-3 localizes broadly to somatic and germline nuclei where it appears enriched on chromatin, including within the pachytene region of the germline (Fig. 3E), a localization consistent with a possible role in piRNA expression.

### CK2 directly Phosphorylates USTC factor TOFU-4

The USTC is thought to bind the promoters of piRNA genes to drive their expression in the *C. elegans* germline. TOFU-4 is a component of the USTC and KIN-3 was identified as a yeast two-hybrid (Y2H) binding partner of TOFU-4 ^10^. We therefore asked if CK2 interacts with TOFU-4. We found that KIN-3 co-immunoprecipitated TOFU-4 (Fig. 4A). Since CK2 is a conserved serine/threonine kinase, we examined whether CK2 can phosphorylate TOFU-4 *in vitro*. TOFU-4, KIN-3 and KIN-10 proteins were purified from *E.coli*, and *in vitro* kinase assays were performed. Incubation of KIN-3/KIN-10 (CK2 complex) with TOFU-4 resulted in a strong phosphorylation signal in *in vitro* phosphorylation assays (Fig. 4B), but KIN-3/KIN-10 failed to phosphorylate the Glutathione S-transferase (GST) protein (Fig. 4D). Thus, CK2 directly and specifically phosphorylates TOFU-4. Phosphorylation was dependent on CK2 activity, as inhibition of CK2 activity by addition of 4,5,6,7-tetrabromobenzotriazole (TBB), a CK2 inhibitor, reduced phosphorylation (Fig. 4B). A *C. elegans* phosphoproteome dataset ^26^ identified three phosphorylation sites in TOFU-4, each containing a consensus CK2 phosphorylation motif ([S/T]-X-X-[D/E]) (Fig. 4C). We found that synthetic TOFU-4 peptides containing either serine 131 (S131) or serine 229 (S229), were phosphorylated by CK2 *in vitro* (Fig. 4D). While a peptide containing serine 92(S92) was not phosphorylated. Using AlphaFold, we predicted the structure of TOFU-4 and discovered that S131 and S229 are in a disordered region and relatively conserved in nematodes (Fig. S3A, B). Consistent with the idea that S131 and S229 are the predominant phosphorylation sites, a mutation of both the S131 and S229 to alanine in a bacterially-expressed full-length TOFU-4 protein greatly reduced phosphorylation of TOFU-4 in an *in vitro* CK2 kinase assay (Fig. 4B).

**Figure 4.**
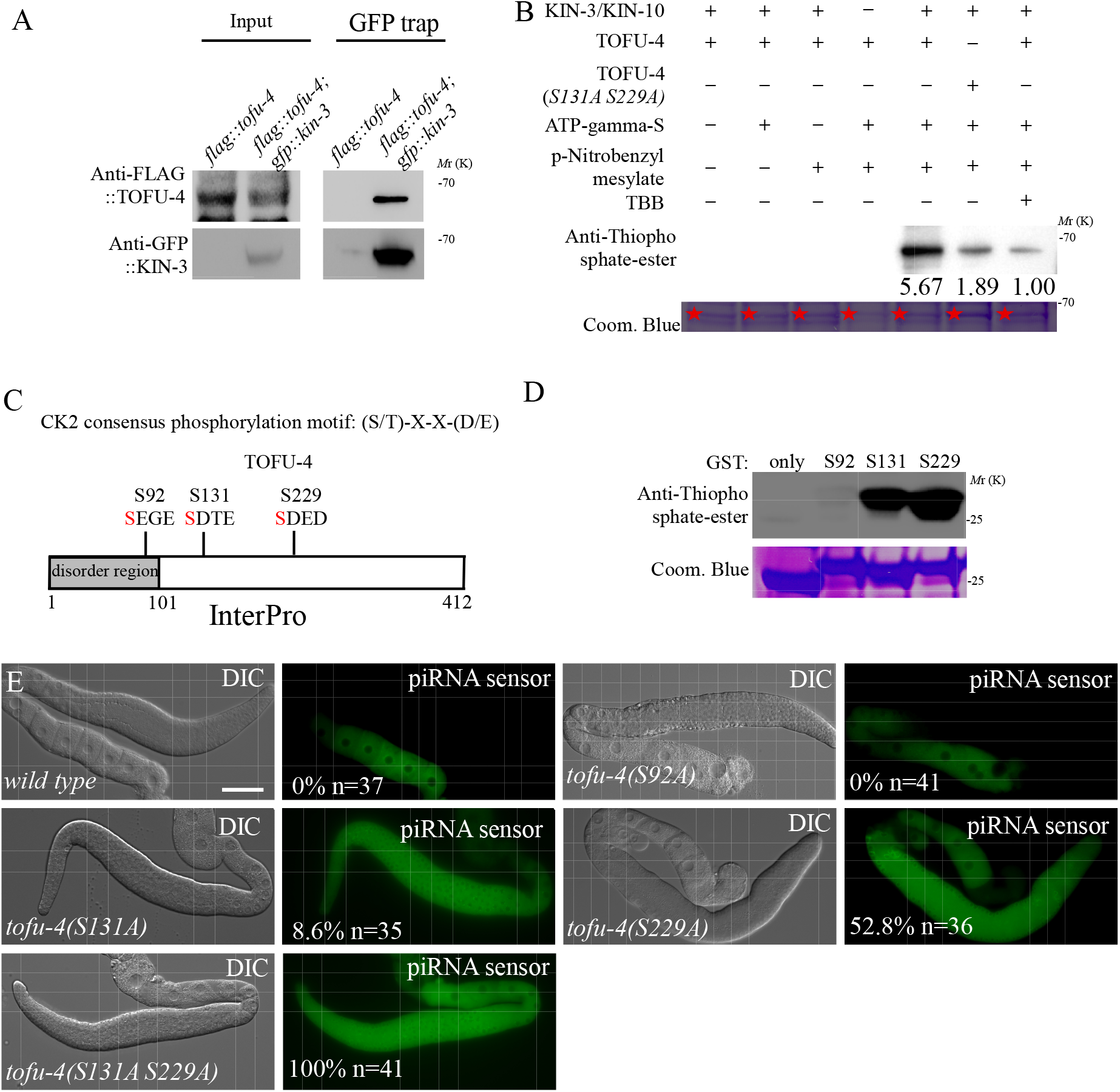
CK2 Phosphorylates USTC factor TOFU-4. **(A)** In co-IP assays, endogenous GFP::KIN-3 specifically immunoprecipitated endogenous FLAG::TOFU-4. **(B)** Phosphorylation of TOFU-4 by the CK2 complex in an *in vitro* phosphorylation assay. Phosphorylation was detected by an antibody that specifically detects the thiophosphate-ester. The purified TOFU-4 (indicated by asterisks) was used in the assay. Coom. Blue, Coomassie staining. Quantifications of the phosphorylation are also shown. The phosphorylation level in the last lane is set to 1.0. **(C)** Three phosphorylation sites were identified in TOFU-4, all of which are located within CK2 recognition motifs from a *C. elegans* phosphoproteome dataset. **(D)** *In vitro* phosphorylation assay showed that Serine 131 and Serine 229 were phosphorylated by CK2. TOFU-4 peptides composed of 20 residues flanking each putative CK2 phosphorylation site were used in the assay. Coom. Blue, Coomassie staining. **(E)** The GFP::CSR-1 transgene is silenced in wild type worms, while in *tofu-4(S131A), tofu-4(S229A)* and *tofu-4(S131A S229A)* mutants worms, GFP::CSR-1 is expressed. Scale bars: 50 μm for E

To determine whether CK2 phosphorylation of TOFU-4 promotes piRNA silencing, we used CRISPR to introduce phosphoacceptor mutants in TOFU-4. As expected, based on the lack of phosphorylation in *in vitro* kinase assays, S92A mutants had no defect in piRNA silencing (Fig. 4E). The S131A and S229A lesions, on the other hand, each caused partial de-silencing individually, and complete desilencing when combined in *tofu-4(S131A S229A)* double mutant worms (Fig. 4E). Additionally, we generated a phosphomimic mutant of TOFU-4, denoted as *tofu-4(S131D S229D)*, in which the potential phospho-acceptor residues were replaced with aspartic acid. Aspartic acid is a negatively charged residue that can sometimes mimic the phosphorylated state of a protein. Surprisingly, in this mutant, we observed a significant desilencing of the piRNA sensor (Fig. S3C), this suggests that the phosphomimic mutant TOFU-4(S131D S229D) is unable to effectively mimic the phosphorylated state of TOFU-4. Taken together these results suggest that phosphorylation of serines 131 and 229 promotes TOFU-4 function.

### CK2 Phosphorylation of TOFU-4 promotes USTC assembly

To explore the stability and protein interactions of the TOFU-4 phospho-acceptor mutants we performed Western blot and Co-IP studies. PRDE-1 and TOFU-4 protein levels appeared wild type, as measured by Western blotting, in *tofu-4(S131A S229A)* mutants (Fig. 5A). To further characterize the effect of these mutations on USTC assembly, we examined their effects on TOFU-4/PRDE-1 protein-protein interactions *in vivo*. We found that PRDE-1 co-precipitated with wild type TOFU-4 (Fig. 5B). However, TOFU-4(S131A S229A) exhibited decreased levels of co-precipitation.

**Figure 5.**
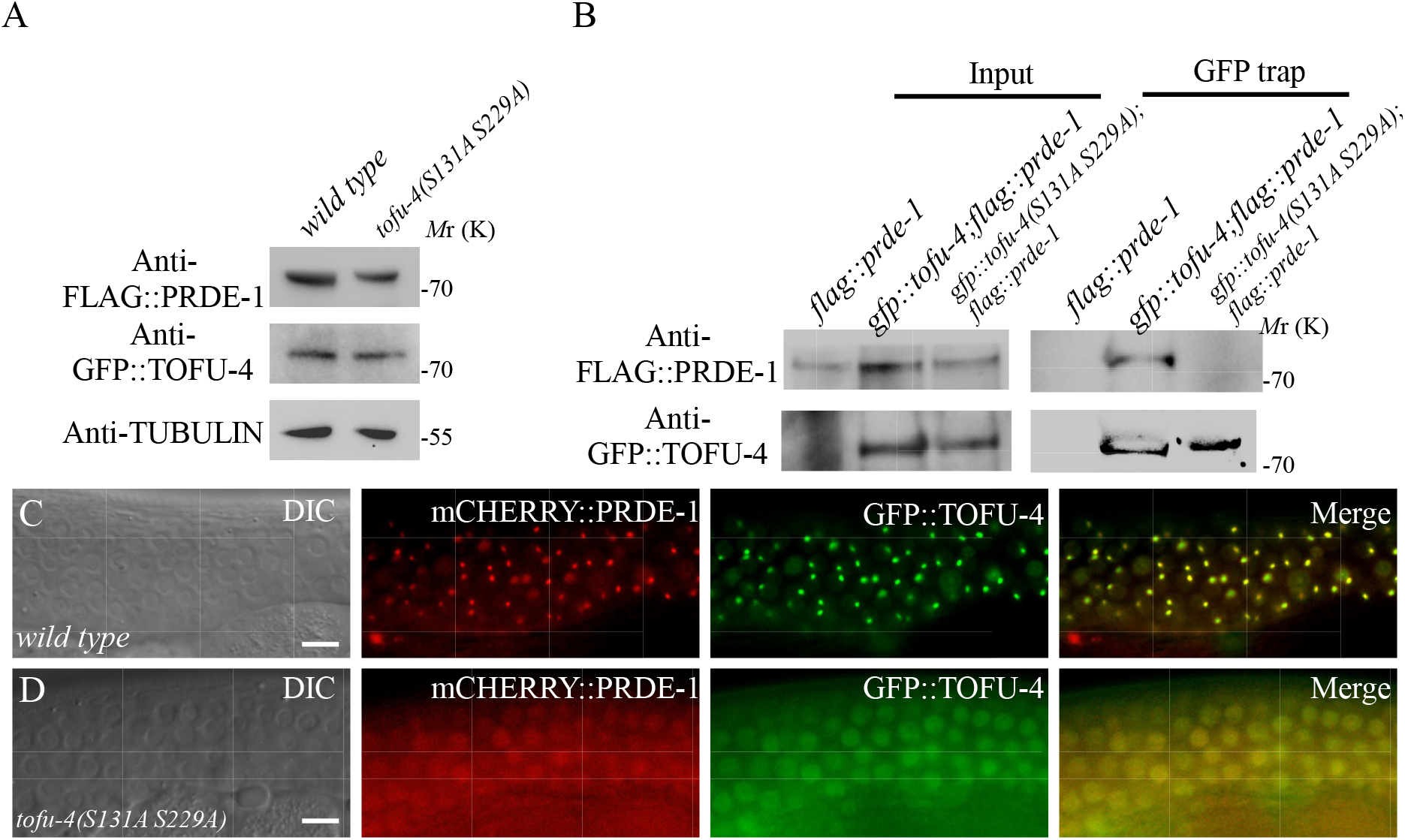
CK2 mediated Phosphorylation of TOFU-4 promotes USTC assembly. **(A)** Protein levels of GFP::TOFU-4 and FLAG::PRDE-1 in wild type and in *tofu-4(S131A S229A)* mutants. **(B)** Western blot analysis of FLAG::PRDE-1 in GFP::TOFU-4 immunoprecipitants from wild type and *tofu-4(S131A S229A)* mutants. **(C, D)** Compared to the wild type worms (C), USTC factors GFP::TOFU-4 and mCHERRY::PRDE-1 fail to form nuclear foci in the germline in *tofu-4(S131A S229A)* mutants worms (D). Scale bars: 20 μm (C and D)

In wild type worms, PRDE-1 and TOFU-4 form subnuclear foci in germline cells and colocalize with each other (Fig. 5C). We therefore examined whether and how the putative phospho-acceptor lesions affected their localization. Introduction of TOFU-4(S131A S229A) in an otherwise wild type strain caused mCHERRY::PRDE-1 and GFP::TOFU-4 to no longer co-localize and to instead become diffusely localized in nuclei (Fig. 5D). Taken together these data suggest that phosphorylation of TOFU-4 by CK2 promotes the co-assembly of TOFU-4 with PRDE-1 and their co-localization at sites of piRNA transcription in *C. elegans*.

### Reduced CK2 activity underlies piRNA mediated gene silencing defects in the aging process

Lastly, we monitored CK2 localization and activity in the aging process since a previous study showed that downregulation of CK2 activity promotes the expression of age-related biomarkers in *C. elegans* ^27^. We first examined the localization of KIN-3::GFP, which showed widespread expression, with a stronger presence in the nucleus. However, we found that during aging process, the distinct pattern diminished (Fig. 6A). Subsequently, we investigated if the catalytic activity of CK2 in *C. elegans* was influenced in the aging process. The phosphotransferase activity of CK2 exhibited a 70% decline in worms at day 7 as compared to those at day 1 (Fig. 6B). To further investigate the consequences of reduced CK2 activity, we examined the localization of USTC components. Consistent with our hypothesis, we observed the disappearance of GFP::TOFU-4 and mCHERRY::PRDE-1 foci (Fig. 6C, D). We also investigated the localization of GFP::PRG-1 and noted the loss of its perinuclear pattern(Fig. 6E). Moreover, western blotting analysis revealed a substantial reduction in the band corresponding to full-length GFP::PRG-1 protein in worms at day 7 (Fig. 6F). Finally, we assessed whether the aging process affected our piRNA reporter and transposable elements. Our findings indicated significant desilencing of the piRNA reporter and notable increases in the *line2h* and *turmoil1* transposons (Fig. 6G-I). Consequently, reduced CK2 activity serves as the underlying cause of piRNA-mediated gene silencing defects during the aging process.

**Figure 6.**
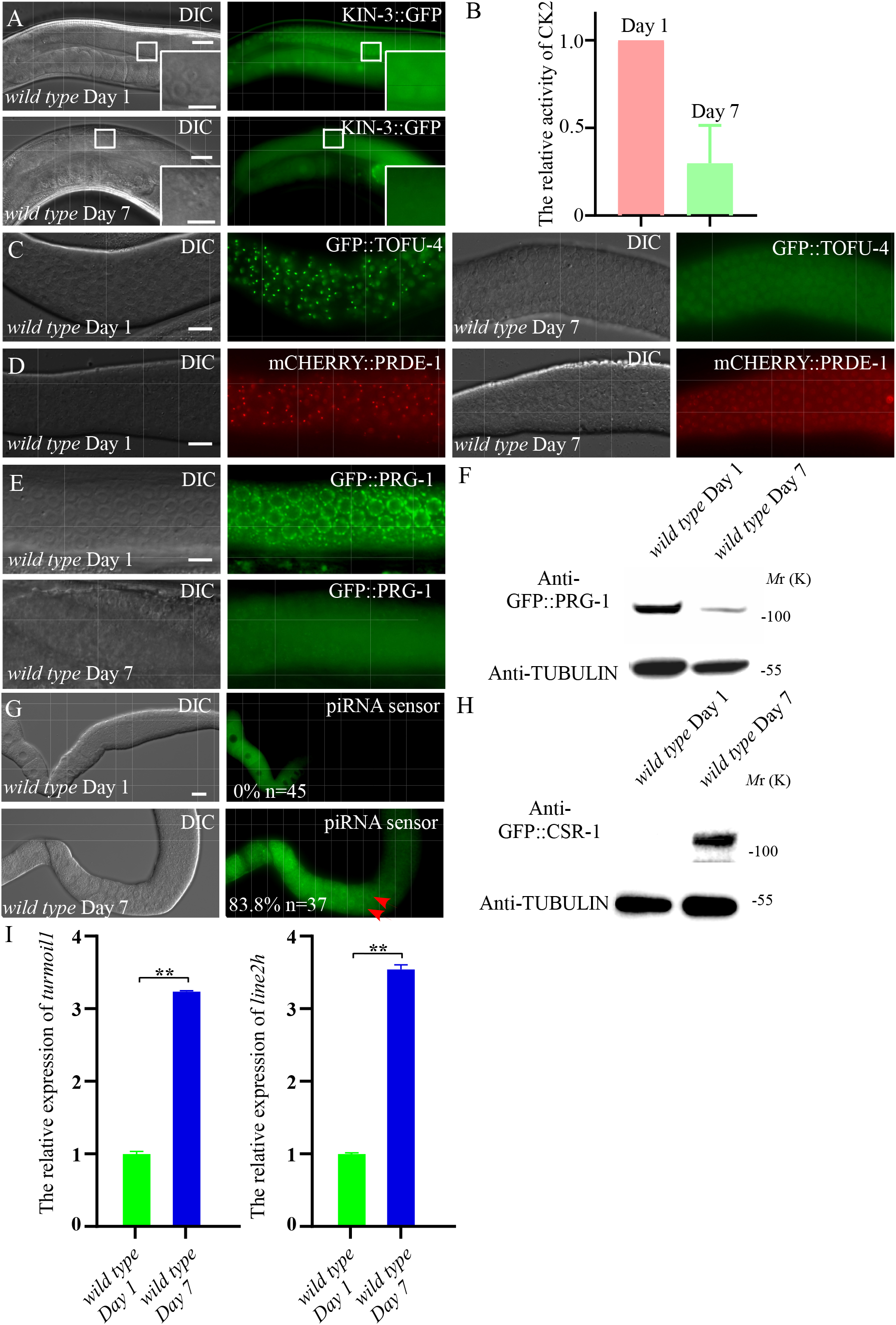
Reduced CK2 activity in the aging process underlies piRNA mediated gene silencing defects. **(A)** Subcellular localization of KIN-3::GFP in wild-type worms at Day 1 and Day 7. **(B)** The activity of CK2 to phosphorylate purified TOFU-4 peptide is decreased in the aging process. Lysates from worms at days 1 and 7 of adulthood were used in *in vitro* phosphorylation assays. **(C, D)** Compared to the wild-type worms at Day 1, the nuclear foci of USTC factors GFP::TOFU-4(C) and mCHERRY::PRDE-1(D) disappear. **(E)** GFP::PRG-1 becomes diffused in the wild-type worms at Day 7 compared to those of wild-type worms at Day 1. **(F)** Immunoblotting assays revealed that protein levels of GFP::PRG-1 are decreased in extracts of wild-type worms at Day 7 compared to those of wild-type worms at Day 1. **(G)** Percentage of worms with expressed piRNA sensors in wide-type worms at Day 1 and wild-type worms at Day 7. n. Total number of animals scored. **(H)** Western results of GFP::CSR-1 protein levels in wide-type worms at Day 1 and wild-type worms at Day 7. **(I)** qRT–PCR of *turmoil1* and *line2h* transposon levels in wild-type worms at Day 1 and wild-type worms at Day 7 relative to act-3 mRNA are displayed. mRNA level in wild-type worms is set to 1.0. Data is shown as mean ± SD. **p < 0.01. Scale bars: 20 μm (A and C-E), 50 μm for G and 10 μm (insets in A)

## Discussion

### A sensitized reporter to screen mutants involved in piRNA mediated gene silencing

In diverse metazoans piRNAs play maintain genome integrity by silencing transposons and have also been implicated in the transgenerational regulation of gene expression^28^. While significant progress has been made in understanding piRNA biogenesis and function, many gaps in understanding persist, especially around elements of the pathway that are shared by other essential germline pathways. Here we constructed a sensitized reporter strain and used it to screen a library of 1000 known essential *C. elegans* genes for RNAi-induced loss of piRNA silencing phenotypes. This screen identified 39 positive hits including previously identified genes required for piRNA biogenesis and function, including components of chromatin remodeling complexes, splicing factors, nuclear export factors, and transcription factors. Our screen identified about 20 genes not previously known to function in the piRNA pathway and here we have focused on characterizing one of these, *kin-10,* which along with *kin-3* comprise components of the CK2 kinase complex.

### TOFU-4 phosphorylation by CK2 promotes piRNA mediated gene silencing

The biogenesis of piRNAs involves a complex series of events, including transcription, processing, and loading into piRNA effector complexes. Previous studies have shown that USTC containing PRDE-1, SNPC-4, TOFU-4, and TOFU-5 is required for the expression of piRNA precursors ^10^. Yet, little is known about how piRNA biogenesis is regulated. Here we have shown that the *C. elegans* casein kinase II, whose major subunits are encoded by the *kin-3* and *kin-10* genes, is required for piRNA mediated silencing. Mechanistically, our analyses place CK2 function upstream of USTC. We also found that TOFU-4 is a direct CK2 substrate. Phosphorylation of TOFU-4 by CK2 influences USTC assembly. In the CK2 mutants, USTC did not assemble, and piRNA biogenesis was defective (Fig. 7). Our study demonstrates that posttranslational modification of the piRNA biogenesis machinery is important for the piRNA pathway.

**Figure 7.**
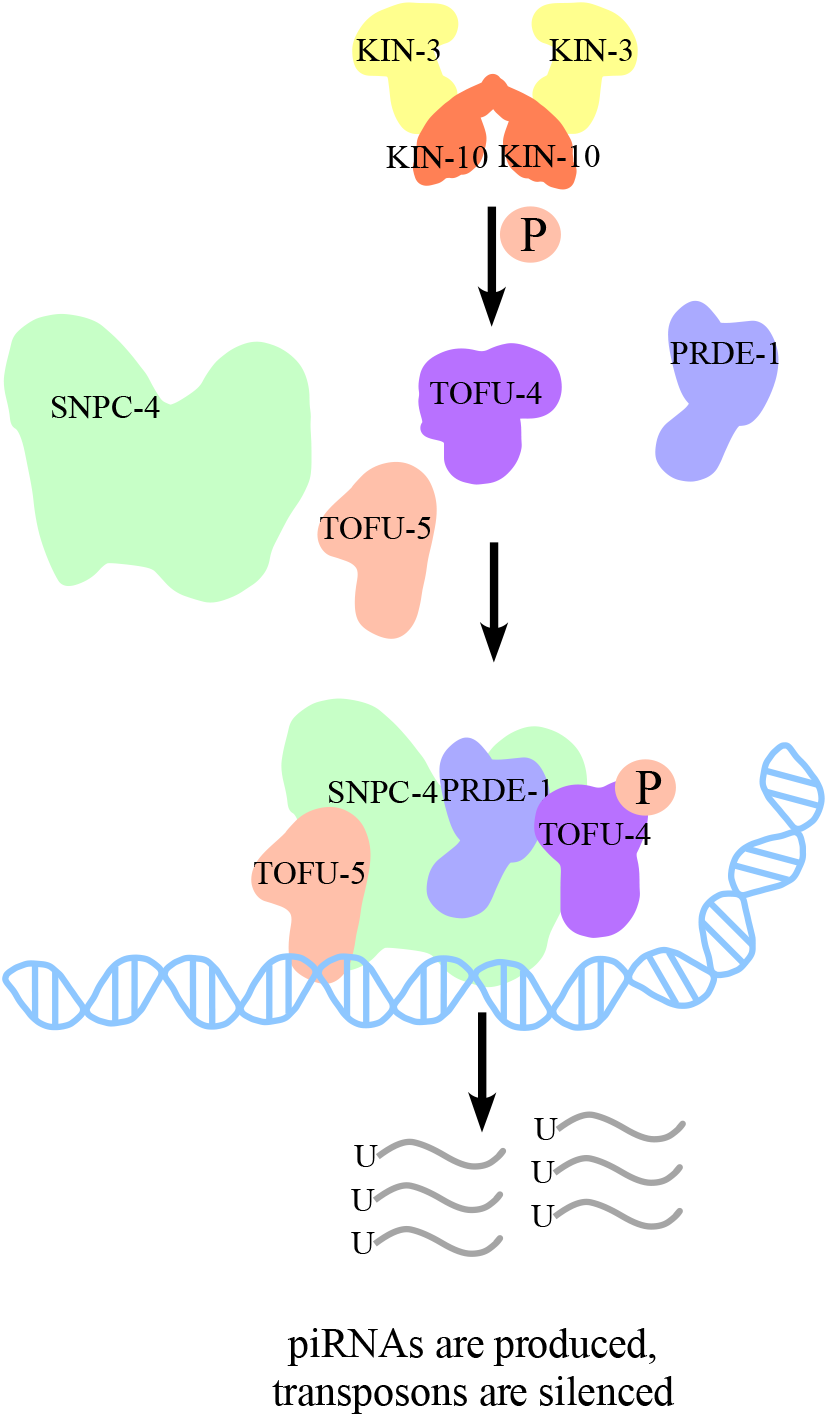
CK2 mediated phosphorylation of TOFU-4 promotes piRNA mediated silencing. A proposed model representing the regulation of phosphorylation of TOFU-4 by CK2 promotes USTC assembly and piRNA biogenesis in the germline. The production of piRNAs silence the foreign transgenes and transposons.

Our findings suggest that phosphorylation of TOFU-4 promotes its association with PRDE-1 and the co-assembly of the two proteins on piRNA cluster genes where along with other components of the USTC they promote piRNA transcription. In *S. pombe,* CK2 directed phosphorylation of a component of the shelterin telomere protein complex, Rap1, promotes interactions with factors that regulate proper telomere tethering to the inner nuclear envelop and the formation of the silenced chromatin structure at chromosome ends ^16^. It will be interesting to learn whether CK2 phosphorylation of TOFU-4 promotes the subnuclear localization of the huge, megabase-scale, clusters of piRNA genes to coordinate piRNA transcription with piRNA export and processing.

### Aging and piRNA mediated gene silencing

During the aging process, loss of heterochromatin is a consistent trend in species ranging from humans to fungi ^29^. How and why heterochromatin is lost in the aging process is not known and the mechanisms are complex and generally cell-type specific. Since piRNA pathways promote the maintenance and installation of heterochromatin in diverse animal germlines it is possible that CK2 promotes heterochromatin maintenance, at least in part, through the maintenance of piRNA activity. Considering that heterochromatin maintenance is implicated in lifespan regulation^29^, further work is needed to explore all the possible relationships between CK2 activity, piRNAs and lifespan.

In summary, our work has uncovered a role for CK2 in the regulation of piRNA biogenesis in *C. elegans* and provides a starting point for a more detailed examination of the role of post-translational modifications in piRNA biogenesis in the aging process.

## Material and methods

### *C. elegans* Strains and Bacterial Strains

All the strains in this study were derived from Bristol N2 and cultured at 20 °C on Nematode Growth Media (NGM) unless otherwise indicated^30^. The strains used in this study are listed in Table S1.

### RNAi screen

RNAi screen was performed against all 947 genes in the embryo lethal subset of the *C. elegans* RNAi collection (Ahringer). Synchronized L1 animals of the reporter strain were plated onto RNAi feeding plates with about 100 worms per plate and were grown at 20°C. The desilencing phenotype was scored when the worms grew to L4 or young adult stage.

### CRISPR/Cas9 genome editing

For CRISPR/Cas9 experiments, Cas9 ribonucleoprotein (RNP) editing was used to generate the CRISPER lines ^31^. Cas9 genome editing mixture containing Cas9 protein (0.5μl of 10µg/µl), two crRNAs (each 1.4μl of 0.4μg/μl), annealed PCR donor (25ng/μl), and PRF4::*rol-6(su1006)* plasmid (0.8μl of 500ng/μl) was incubated at 37°C for 15mins before injecting animals. Guide RNA sequences and donors used in this study are listed in Table S2.

### RNAi inactivation experiments in *C. elegans*

For RNAi inactivation, single-stranded RNAs (ssRNAs) were transcribed from SP6-flanked PCR templates. ssRNAs were then annealed and injected into animals carrying various reporters. Injected animals were placed on fresh plates overnight. The progeny was analyzed.

### Auxin treatment

The auxin-inducible degron system was as described ^32^. The degron-tagged L1 larvae were plated on NGM plates with 500 μM indole-3-acetic acid (IAA; Alfa Aesar, 10171307). Worms were placed on IAA plates and collected at young adult stage for further analysis.

### Co-IP and western blotting

Synchronous adult worms were collected and washed three times with M9 buffer. The worms were then homogenized in a FastPrep-24 benchtop homogenizer (MP Biomedicals). Worm extracts were centrifuged at 14000 × *g* for 15 mins at 4°C. The worm extracts were incubated with GFP trap beads for 2 hrs at 4°C on a rotating shaker. The beads were washed three times and subjected to SDS-PAGE. Signals were detected using corresponding primary and secondary antibodies.

### Small RNA cloning and data analysis

The small RNA cloning was conducted as the previous paper ^22^. Total RNAs were extracted by Trizol (Sigma Alrich) and small RNAs were enriched by an mir-Vana miRNA isolation kit (Thermo Scientific). Samples were pretreated with homemade PIR-1 or 5’ Pyrophosphohydrolase (RppH, NEB). The small RNAs were then ligated to a 3’ adaptor (5’ rAppAGATCGGAAGAGCACACGTCTGAACTCCAGTCA/3ddC/3’; IDT) by T4 RNA ligase 2(NEB). The 5’ adaptor containing 6 nt barcode was ligated using T4 RNA ligase 1. The ligated products were reverse transcribed using SuperScript III (Thermo Fisher Scientific). The cDNAs were amplified by PCR and the libraries were sequenced using the HiSeq systems (Illumina) platform at the UMass Medical School Deep Sequencing Core Facility.

The small RNA sequencing data were analyzed as described ^22^. Briefly, adaptors were trimmed from fastq files. Reads were trimmed to retain only 18-30 nts. The reads were mapped to piRNAs or piRNAs precursors with an arbitrary length of overhanging sequence either side of the 21U sequence using the exact matches. Reads were normalized to the total reads mapped to the genome.

### Protein expression and purification

All genes were amplified by PCR and cloned into the pET.28a vector to produce His6-tag-fused recombinant proteins, or the pGEX-6P-1 vector to produce GST-tag-fused recombinant proteins. All recombinant proteins used in this study were expressed in *E. coli* BL21-CodonPlus (DE3) induced by 0.3 mM IPTG for 16 h at 20°C and collected by sedimentation.

To purify His-KIN-3, His-KIN-10, His-TOFU-4 and GST-tagged TOFU-4 fragments, the *E. coli* cells were resuspended in binding buffer (50 mM Tris-Cl pH 7.9, 500 mM NaCl and 10 mM imidazole), lysed with a high-pressure homogenizer and sedimented at 18000 rpm for 30 min at 4°C. The supernatant lysates were purified on Ni-NTA agarose beads (for His6-tagged proteins, Qiagen) or Glutathione High Capacity Magnetic Agarode Beads (for GST-tagged proteins, SIGMA). After 2 extensive washing with binding buffer, the proteins were eluted with His6 elution buffer (50 mM Tris-Cl pH 7.9, 500 mM NaCl and 500 mM imidazole, for His6-tagged proteins) or GST elution buffer (50 mM Tris-Cl pH 7.9, 500 mM NaCl and 10 mM GSH, for GST-tagged proteins). All the eluted proteins were concentrated by centrifugal filtrations (Millipore), loaded onto desalting columns (GE Healthcare), then eluted with PBS buffer (140 mM NaCl, 2.7 mM KCl, 10 mM Na_2_HPO_4_, and 1.8 mM KH_2_PO_4_). The eluted proteins were stored in aliquots at −80 °C.

### *In vitro* phosphorylation assays using ATP-γS

The *in vitro* phosphorylation assays using ATP-γS was conducted as the previous paper ^33^. To detect whether a protein is directly phosphorylated by CK2 complex, the His-KIN-3 and His-KIN-10 proteins purified from *E. coli* were used for the kinase assay. Recombinant His-TOFU-4, GST, GST-TOFU-4(S92), GST-TOFU-4(S131) and GST-TOFU-4(S229) proteins purified from *E. coli* were used as substrates. The reaction mixtures containing 500 μM ATP-γS were incubated for 1 h at 30 °C in kinase assay buffer (30 mM HEPES, 50 mM potassium acetate, 5 mM MgCl_2_). For detection by anti-thiophosphate-ester antibody, the reaction mixtures were further supplemented with 2.5 mM *p*-nitrobenzyl mesylate (PNBM). The alkylating reactions were allowed to proceed for 1 h at 25 °C. The reaction systems were terminated with SDS sample buffer and boiled before analysis by SDS-PAGE.

To detect CK2 activity in worms, the assay was conducted as described previously^27^. Briefly, worms were lysed, the protein concentration in supernatant was measured and adjusted. The *in vitro* phosphorylation assay was conducted using ATP-γS. GST-TOFU-4(S229) proteins purified from *E. coli* were used as substrates The reactions were initiated by adding worm lysates and incubated for 30 mins.

## Data Availability

The small RNA sequencing data are available from GEO.

## Statistical analysis

GraphPad Prism 7 or Microsoft Excel was used for all statistical analysis. All data are shown as mean ± SD. Unpaired two tailed t tests were performed for statistical analysis. a P-value less than 0.01 was considered extremely significant (**).

For the small RNA seq, two repeats were perfirmed. For other experiments, at least 3 independent repeats were performed.

## Acknowledgements

We thank members of Mello lab for discussions, Caenorhabditis Genetics Center (CGC) for providing strain, Dr. Weifeng Gu for providing the PIR-1 protein for preparing small RNA sequencing libraries. This work was supported by the following grants to C.C.M.: NIH grants (GM058800 and HD078253), C.C.M. is a Howard Hughes Medical Institute Investigator.

## Author contributions

G.M.Z. and C.C.M. designed the experiments. G.M.Z. and Y.H.D performed the experiments. C.W.Z analyzed the sequencing results. G.M.Z. and C.C.M. wrote the manuscript.

## Supplemental Figure legends

**Supplementary Figure 1.**
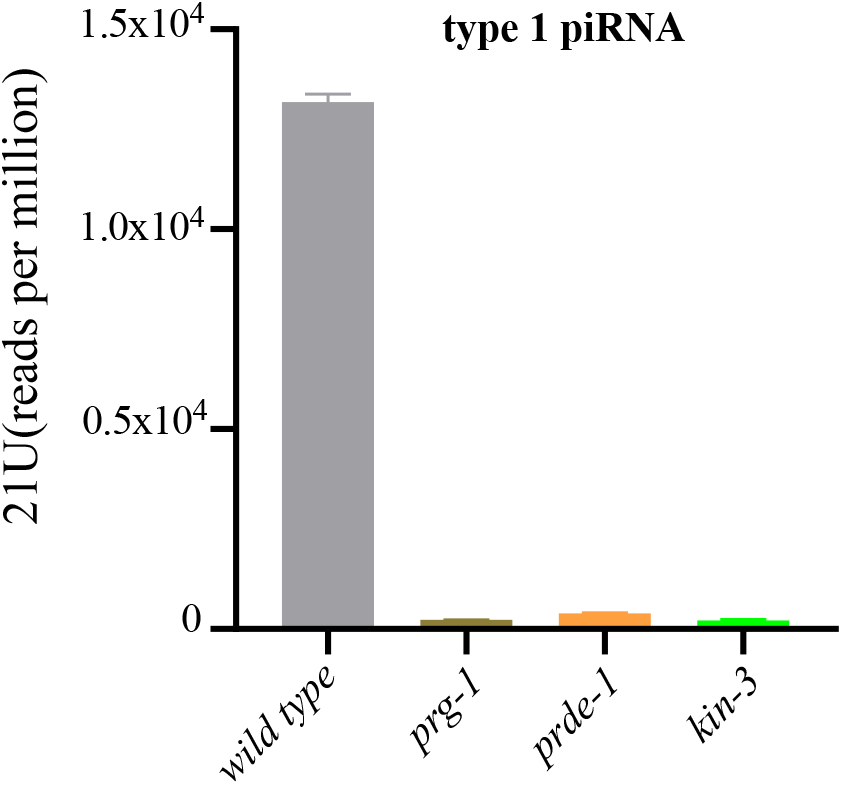
CK2 complex affects type I piRNAs level. Bar diagram displaying type 1 21U-RNA abundance in wild type, *prg-1*, *prde-1*, and *kin-3* mutants.

**Supplementary Figure 2.**
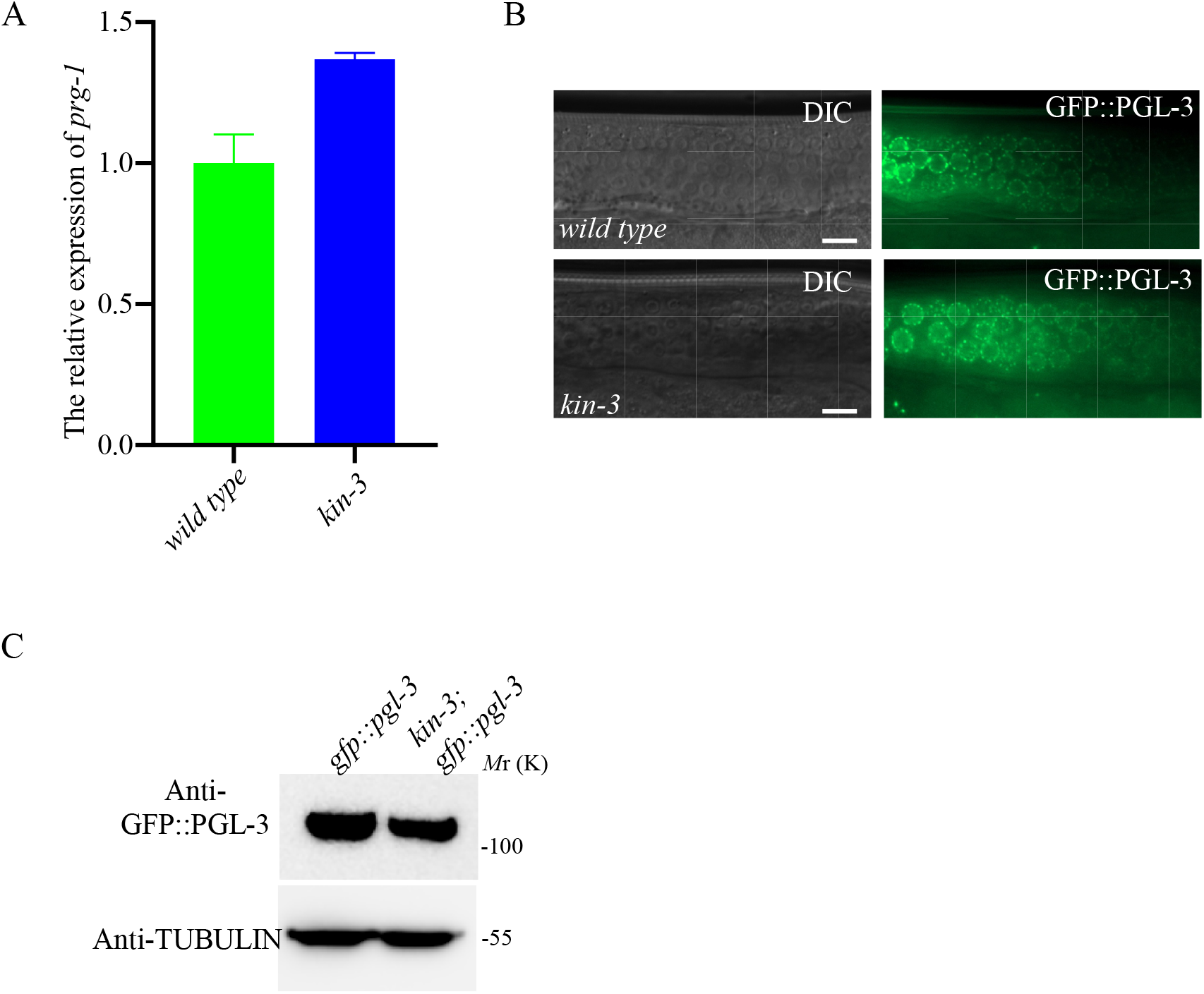
CK2 complex does not affect the expression of GFP::PGL-3. **(A)** qRT–PCR results of *prg-1* levels in wild type worms and in *kin-3* mutants worms relative to *act-3* mRNA are displayed. mRNA level in wild type worms is set to 1.0. Data are shown as mean ± SD. **(B)** The expression of GFP::PGL-3 in the wild type worms is similar to that in *kin-3* mutants. **(C)** Protein levels of GFP::PGL-3 are similar in extracts of *kin-3* mutants compared to those of wild type worms. Scale bars: 10 μm in B

**Supplementary Figure 3.**
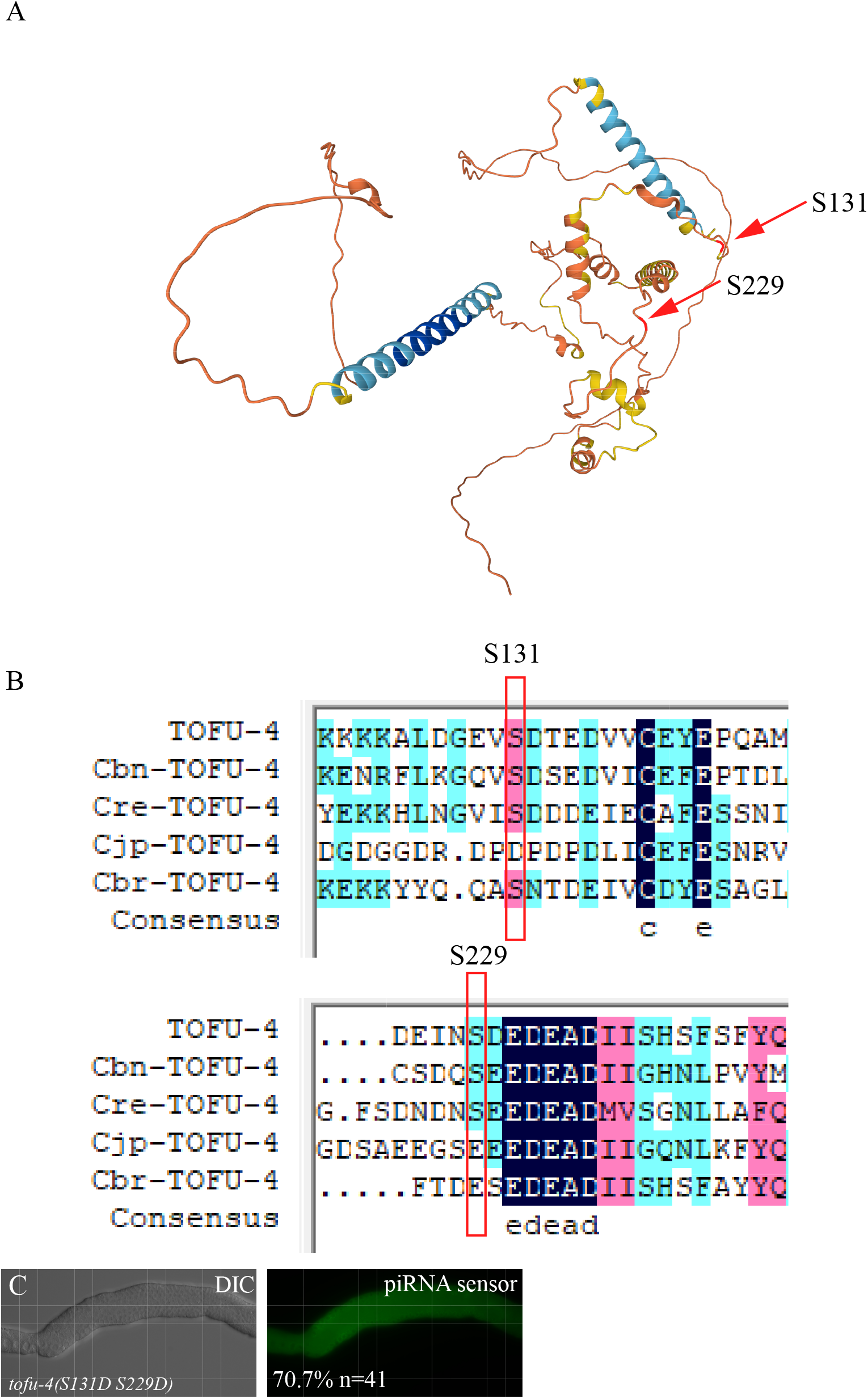
TOFU-4 protein structure and sequence homology. **(A)** Protein structural predictions for TOFU-4 is shown. Protein structure was predicted using the Alphafold (https://alphafold.ebi.ac.uk/). Position of S131 and S229 were indicated by arrowhead. **(B)** Alignment of different nematodes TOFU-4 peptides is shown. Position of S131 and S229 were indicated. **(C)** The GFP::CSR-1 transgene is largely desilenced *tofu-4(S131D S229D)* worms. Scale bars: 50 μm for C

